# RP3V kisspeptin neurons mediate neuroprogesterone induction of the luteinizing hormone surge in female rat

**DOI:** 10.1101/700435

**Authors:** Lauren K. Delhousay, Timbora Chuon, Melinda Mittleman-Smith, Paul Micevych, Kevin Sinchak

## Abstract

To induce ovulation, neural circuits are sequentially activated by estradiol and progesterone. In female rodents, estradiol-induced neuroprogesterone, synthesized in astrocytes, is essential for the luteinizing hormone (LH) surge and subsequently, ovulation. However, the gonadotropin-releasing hormone (GnRH) neurons do not express the steroid receptors necessary for reproduction: progesterone receptors (PGR) or estrogen receptor-α (ERα). Steroid information is transduced by rostral periventricular (RP3V) kisspeptin neurons that express ERα and PGR and innervate GnRH neurons in the diagonal band of Broca (DBB) and the medial septum. In this study, we tested the hypothesis that estradiol induced neuroprogesterone needed for the LH surge is mediated by kisspeptin. Neuroprogesterone synthesis was inhibited with aminoglutethimide (AGT; s.c.) in 17β-estradiol benzoate (EB)-primed, ovariectomized (ovx) and adrenalectomized (adx) rats. Kisspeptin-10 (20 nmol/µl) was infused into the DBB, trunk blood was collected 53 hours post-EB injection, and serum LH levels were analyzed by ELISA. AGT inhibition of neuroprogesterone synthesis blocked the EB-induced LH surge. Subsequent treatment with either progesterone or DBB kisspeptin-10 infusion restored the LH surge. Kisspeptin restored the LH surge, which was blocked by DBB infusion of kisspeptin receptor (GPR54) antagonist (kisspeptin-234). Finally, knockdown of kisspeptin protein levels in the RP3V with kisspeptin antisense oligodeoxynucleotide (ODN) significantly lowered LH levels in EB-primed rats compared to scrambled ODN, demonstrating the importance of endogenous RP3V kisspeptin for the LH surge. These results support the hypothesis that neuroprogesterone induces both kisspeptin release from RP3V neurons impacting the LH surge.

## INTRODUCTION

Ovulation, a critical event in female reproduction, is synchronized with the cyclic development of the uterine lining and regulation of reproductive behavior to increase the likelihood that copulation will result in fertilization of the ovulated egg and ultimately, pregnancy. In rats, these events are regulated by hormones released by the hypothalamic-pituitary-gonadal axis. Maturation of ovarian follicles initially causes a slow rise in circulating estradiol levels during diestrus I and II, which primes the luteinizing hormone (LH) surge regulating neurocircuitry by inducing progesterone receptor (PGR) expression and gonadotropin releasing hormone (GnRH) synthesis (Chappell & Levine, 2000). On proestrus, estradiol levels rise rapidly to peak in the afternoon inducing estrogen positive feedback, resulting in the preovultatory LH surge that stimulates ovulation and follicular luteinization (Chazal, Faudon, Gogan, & Laplante, 1974). Over the years, it has become clear that in addition to estradiol, progesterone is also required (DePaolo & Barraclough, 1979; P. Micevych et al., 2003; P.E. Micevych et al., 2007). The proestrous levels of estradiol induce hypothalamic astrocytes to synthesize neuroprogesterone that is essential for the LH surge (P. Micevych et al., 2003; P. E. Micevych et al., 2007; P.E. Micevych et al., 2007; P. E. Micevych & Sinchak, 2011; Figure 1).

**FIGURE 1.**
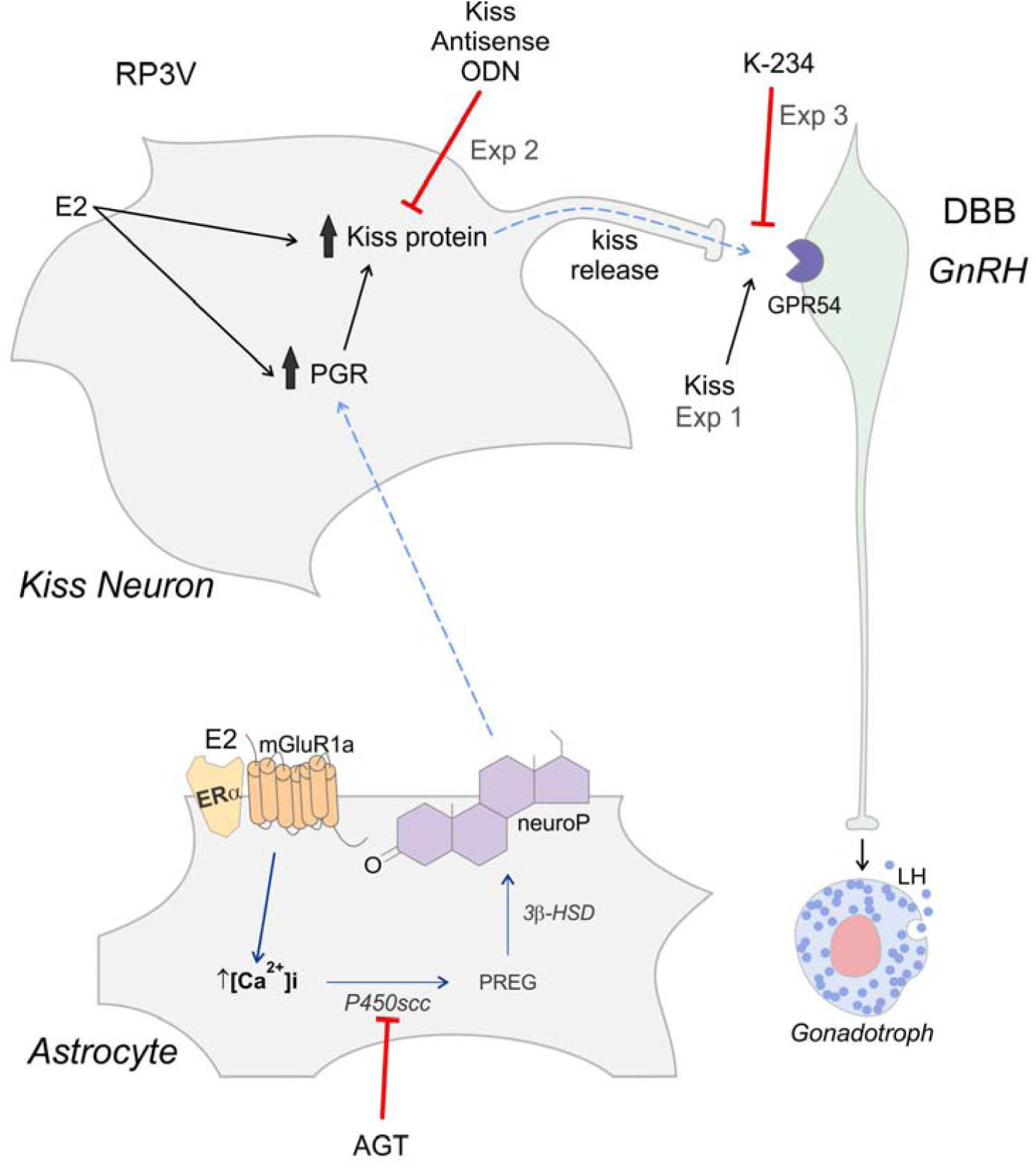
Schema of hypothesized kisspeptin mediation of neuroprogesterone (neuroP) induction of the LH surge in the rat. During the negative feedback of the estrous cycle, estradiol (E2) induces progesterone receptors (PGR) and kisspeptin (Kiss) expression in Kiss neurons of the RP3V. During positive feedback, estradiol acts through a membrane associated estrogen receptor-α that complexes with and signals through metabotropic glutamate receptor-1a (mGluR1a). This produces calcium signaling (Ca^2+^) that initiates the synthesis of neuroP in hypothalamic astrocytes in the RP3V region. This neuroP then acts through E2 induced PGR in Kiss neurons to stimulate the release of Kiss into the diagonal band of Broca. Here Kiss acts through the kisspeptin receptor (GPR54) to excite gonadotropin releasing hormone (GnRH) neurons, releasing GnRH responsible for inducing the LH surge. In experiment 1, neuroprogesterone synthesis was blocked in estradiol primed rats and kisspeptin was infused into the DBB to restore the LH surge to demonstrate that Kiss acts downstream of neuroP to induce the LH surge. In experiment 2, to demonstrate that RP3V Kiss neurons mediate neuroP induction of the LH surge, Kiss expression was reduced by infusion of Kiss antisense oligodeoxynucleotides (ODNs) into the 3V at the level of the RP3V. In experiment 3, to demonstrate that Kiss signals through GPR54 in the DBB to mediate neuroP induction of the LH surge, GPR54 antagonist (K-234) was infused into the DBB in estradiol treated animals.

In ovariectomized and adrenalectomized (ovx/adx) rats, exogenous estradiol treatment alone is sufficient for triggering the preovulatory surge of gonadotropins (Mahesh & Brann, 1998; Mann, Korowitz, Macfarland, & Cost, 1976). It is increasingly clear that the actions of progesterone are required to both induce and attain the full magnitude and duration of the LH surge (Mahesh & Brann, 1998; Mann et al., 1976). Indeed, positive feedback levels of estradiol facilitates neuroprogesterone synthesis by hypothalamic astrocytes through the activation of a membrane estrogen receptor-α (mERα) that complexes with metabotropic glutamate receptor-1a (mGluR1a; Figure 1) to increase intracellular calcium levels and activate a calcium sensitive protein kinase A (PKA) stimulating neuroprogesterone synthesis (Chaban, Lakhter, & Micevych, 2004; Chen, Kuo, Wong, & Micevych, 2014; P.E. Micevych et al., 2007). The neuroprogesterone binds to estradiol-induced PGR in kisspeptin neurons (Mittelman-Smith, Wong, Kathiresan, & Micevych, 2015; Mittelman-Smith, Wong, & Micevych, 2018) to trigger the LH surge. Antagonizing either progesterone signaling or synthesis in estradiol-primed ovx/adx animals prevents induction of the LH surge (Chappell & Levine, 2000; Mann et al., 1976; Nath, Bhakta, & Moudgil, 1992). Thus, the brain is not only a target for steroid hormones of peripheral origin, but is also a steroidogenic tissue capable of synthesizing neurosteroids *de novo* from cholesterol (Baulieu, 1981).

GnRH neurons directly responsible for triggering the LH surge do not express ERα) or PGR (Herbison & Theodosis, 1992; Shivers, Harlan, Morrell, & Pfaff, 1983). Therefore, estradiol-induced neuroprogesterone must signal through PGR on an intermediate neuron that is stimulatory to GnRH neurons, to trigger the LH surge. Kisspeptin neurons in the periventricular and anteroventral periventricular (AVPV) nuclei, which make up the rostral periventricular region of the third ventricle (RP3V) meet both of these criteria. RP3V kisspeptin neurons express both ERα and PGR (Clarkson, d’Anglemont de Tassigny, Moreno, Colledge, & Herbison, 2008; Herbison & Theodosis, 1992; Lehman, Merkley, Coolen, & Goodman, 2010; Shivers et al., 1983; Smith, Cunningham, Rissman, Clifton, & Steiner, 2005). Sub-populations of RP3V kisspeptin neurons project onto GnRH neurons in the diagonal band of Broca (DBB) and potently activate them to induce the secretion of LH (Clarkson & Herbison, 2006; Han et al., 2005; Liu, Lee, & Herbison, 2008; Wintermantel et al., 2006). The importance of PGR in kisspeptin neurons for the LH surge was recently demonstrated in female mice with knockout of PGR specifically in kisspeptin neurons (KissPRKOs; Stephens et al., 2015). Compared to wildtype mice, the estradiol-induced LH surge and RP3V c-Fos induction associated with the LH surge was blocked in the KissPRKO mice. Further, these mice had impaired fertility and ovulation. Moreover, immortalized kisspeptin cells derived from adult female hypothalamic neurons (mHypoA51), an *in vitro* model for RP3V kisspeptin neurons, express ERα and PGR (Mittelman-Smith et al., 2015). In these neurons, kisspeptin expression was induced by estradiol and augmented by progesterone (Mittelman-Smith et al., 2018), mimicking the response of RP3V kisspeptin neurons *in vivo*. In the present study, we tested the hypothesis that *in vivo*, estradiol-induced neuroprogesterone acts on RP3V kisspeptin neurons, triggering an LH surge.

## MATERIALS AND METHODS

### Animals

For all experiments, female Long-Evans rats were ovx/adx by the supplier (Charles River, Hollister, CA) at 55-65 days old or 200-225 grams in weight. Animals were housed two per cage in a climate and light controlled room (12/12 light/dark cycle, lights on at 0600 hours) with food and water provided *ad libitum*. Upon arrival, corticosterone (50mg/L; Sigma Aldrich; St. Louis, MO) in saline (0.9%) was given in the drinking water for the duration of the experiment. The California State University, Long Beach and University of California, Los Angeles IACUC Committees approved all procedures.

### Stereotaxic Surgery for Implantation of Guide Cannulae

Animals were deeply anesthetized 13 days (day 13) after ovx/adx surgery (day 0; Figure 2) with isoflurane (Western Medical Supply Inc., Arcadia, CA) injected with an analgesic (Rimadyl 1 mg/ml; Western Medical Supply Inc., Arcadia, CA) and placed into a stereotaxic frame. In Experiments 1 and 3, standard stereotaxic procedures were used to implant bilateral guide cannulae (22 gauge, 8mm below pedestal; Plastics One; Roanoke, VA) directed at the DBB (coordinates from bregma: 0°angle; anterior **+**0.5mm, lateral **±**0.5mm, ventral −4.5mm, −3.3mm tooth bar). In Experiment 2, unilateral guide cannulae (22 gauge, 11 mm below pedestal; Plastics One; Roanoke, VA) were infused into the third ventricle (3V) at the level of the RP3V (coordinates from bregma: 4° angle; anterior +0.1 mm, lateral +0.7 mm, ventral −6.0 mm, −3.3mm tooth bar). Cannulae were anchored to the cranium with dental cement and bone screws. Stylets were inserted to keep the cannulae patent, and protruded 0.5 mm past the guide cannula opening, and covered with a head cap. After surgery, animals were singly housed and received antibiotics orally through the drinking water (0.5 mg/ml trimethoprim and sulfamethoxazole; TMS; Western Medical Supply Inc., Arcadia, CA). Animals were allowed to recover for 7 days prior to steroid treatment and drug infusion beginning on 20 days after ovx/adx surgery (Figure 2).

**FIGURE 2.**
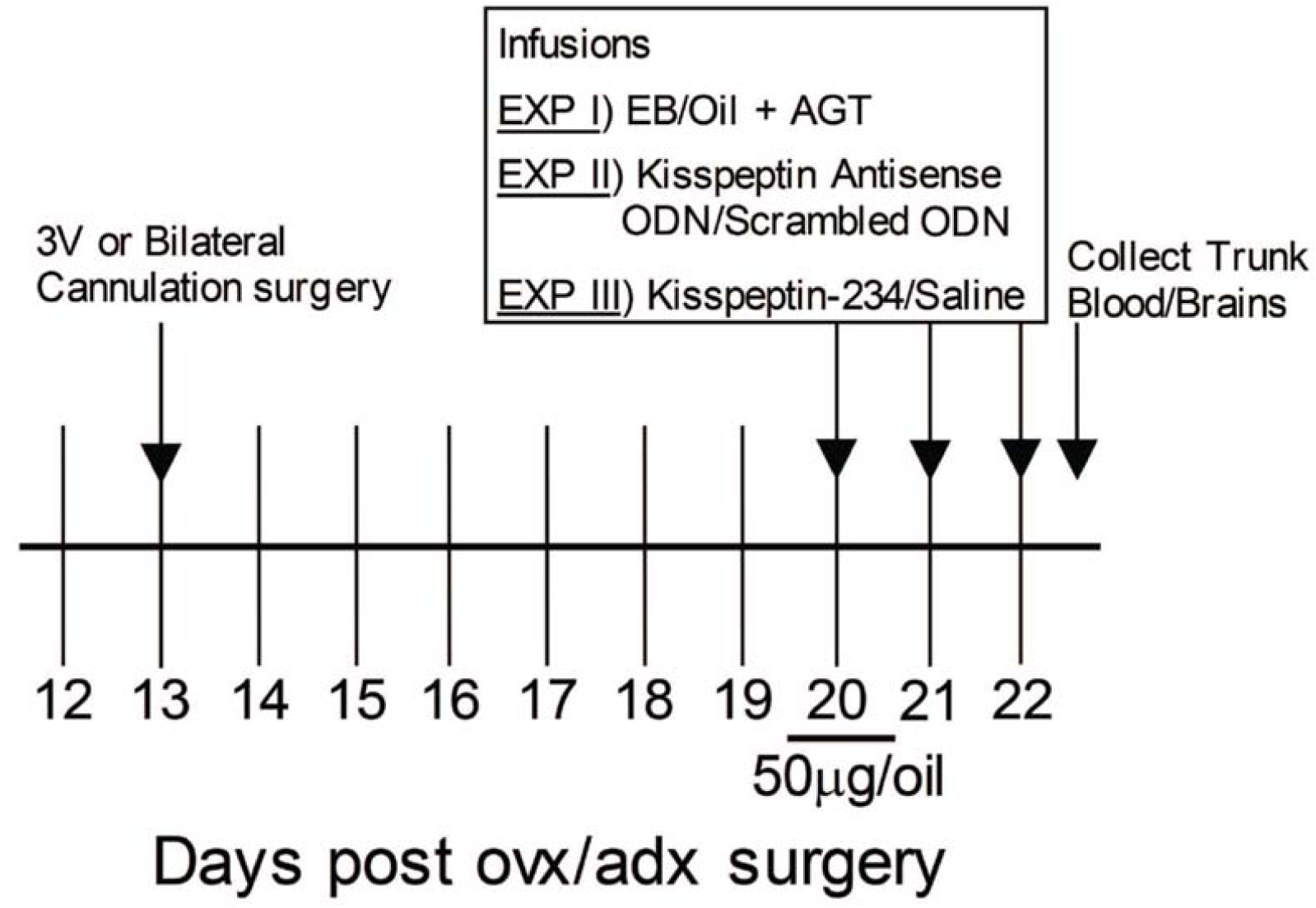
General timeline of experiments. Ovariectomized and adrenalectomized (ovx/adx) Long Evans rats underwent either third ventricular or bilateral cannulation surgeries 13 days post ovx/adx surgeries. They then received either oil or 50 µg 17β-estradiol benzoate (EB; s.c.) 7 days after cannulation surgery (day 20). On this same day, animals received either DBB or 3V infusions, which were repeated on days 21 and 22. Approximately 53 hours post EB treatment, animals were euthanized and trunk blood and brains were collected for serum LH analysis and cannula tract placement, respectively.

### Steroid and Drug Treatments

Animals were treated with either safflower oil (0.1ml; s.c.), 17β-estradiol benzoate (EB; s.c.; 50 µg/0.1ml dissolved in safflower oil) or progesterone (s.c.; 500 μg/0.1 ml dissolved in safflower oil). In Experiments 1 and 2, animals received s.c. injections of The P450 side-chain cleavage inhibitor, aminoglutethimide (AGT), dissolved in DMSO (MP Biomedicals; Santa Ana, CA), or control injections for DMSO were an equivalent volume of DMSO (exact doses given in experimental design).

For kisspeptin knockdown in Experiment 2, scrambled ODN (scODN; control) and kisspeptin asODNs were synthesized as phosphothiorated 20-mers by the supplier (Invitrogen, Grand Island, NY). Three asODNs were designed to be delivered in a cocktail that would target different areas of kisspeptin mRNA to ensure reduction of overall translation into protein (Table 1). Individual kisspeptin asODN and scODNs were dissolved in 0.9% saline at a concentration of 25 µg/1 µl. All 3V infusions of either scODN or kisspeptin asODN cocktails were in a volume of 1 μl.

**Table 1.**
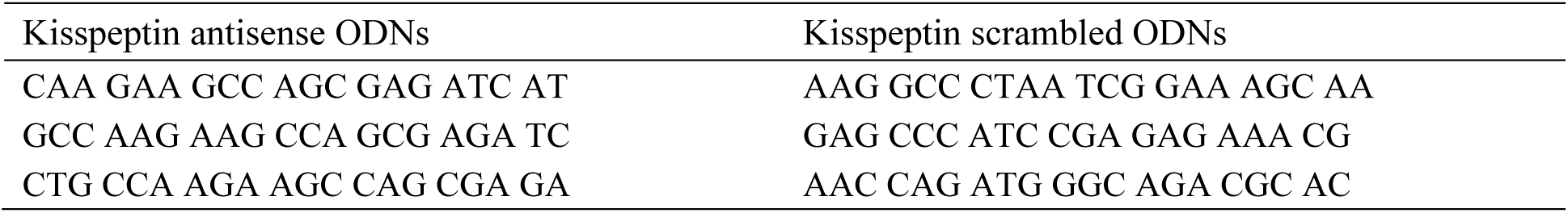
Kisspeptin antisense and scrambled oligodeoxynucleotides used in experiment 2.

In experiments where drugs were site-specifically infused bilaterally, all drugs were dissolved in sterile saline and total volume infused per side was 0.5 μl. All infusions (vehicles, drugs and ODNs) were performed with a sterile microinjector (Plastics One; Roanoke, VA) to a 25 μl Hamilton syringe via thin plastic Tygon™ tubing at a rate of 1 μl per minute by a microliter syringe pump (Stoelting Co., Wood Dale, IL). Microinjectors did not protrude more than 2 mm beyond the opening of the guide cannulae and were left in place for 1 minute after infusion to allow the drug to diffuse away from the injection needle. Stylets were reinserted into the guide cannulae following microinfusion and animals were returned to their home cage until tissue collection.

### Experimental Design

#### Experiment 1

In this experiment, we tested the subhypothesis that kisspeptin acts downstream of neuroprogesterone in the hypothalamic LH surge induction pathway. Briefly, neuroprogesterone synthesis was blocked by treating EB-primed ovx/adx rats with AGT, and GPR54 agonist, kisspeptin-10 (kisspeptin-10; 10 μg/1 µl; Tocris Bioscience; Minneapolis, MN) was administered to the DBB to determine the ability of kisspeptin to rescue the LH surge in the absence of neuroprogesterone. One week before steroid treatment and drug infusions (day 13 after ovx/adx surgery), rats were implanted with a bilateral cannula aimed at the DBB. Starting on day 20 day, all animals were injected with the p450 side chain cleavage inhibitor, AGT, for 3 consecutive days (s.c.; 1 mg/kg on Days 20-21 and 5 mg/kg dissolved in DMSO on Day 22) at 0800 hours. One group of AGT-treated animals (n=7) received a injection of either safflower oil (0.1 ml; s.c.) at 1200 hours on Day 20 and another injection of oil (s.c.) at 1000 hours on Day 22 of the experiment, just prior to bilateral DBB infusion of saline at 1530 hours. A second group of AGT-treated animals (n=7) received EB at 1200 hours on Day 20 and an injection of oil (s.c.) at 1000 hours on Day 22, just prior to DBB infusion of saline at 1530 hours. A third group of AGT-treated animals (n=10) received EB at 1200 hours on Day 20 and an injection of progesterone (500 µg/0.1 ml; s.c.) at 1000 hours on Day 22, just prior to DBB infusion of sterile saline (0.5 µl/per side) at 1530 hours. A fourth group of AGT-treated animals (n=9) received EB at 1200 hours on Day 20 and an injection of safflower oil (0.1 ml; s.c.) at 1000 hours on Day 22, just prior to DBB infusion of kisspeptin (10 nmol/0.5 µl/per side) at 1530 hours. Trunk blood and brains of all animals were collected approximately 53 hours post oil/EB treatment, (1730 hours on Day 22) which is the time of the LH surge in the colony room. Serum LH concentrations were measured using an ELISA kit (Shibayagi via BioVendor; Asheville, NC). Brains were flash-frozen, sectioned at 20 µm and thionin stained for confirmation of cannula placement (Figure 3). Results were analyzed by one-way ANOVA followed by a post-hoc Student-Newman-Keuls (SNK) test.

**FIGURE 3.**
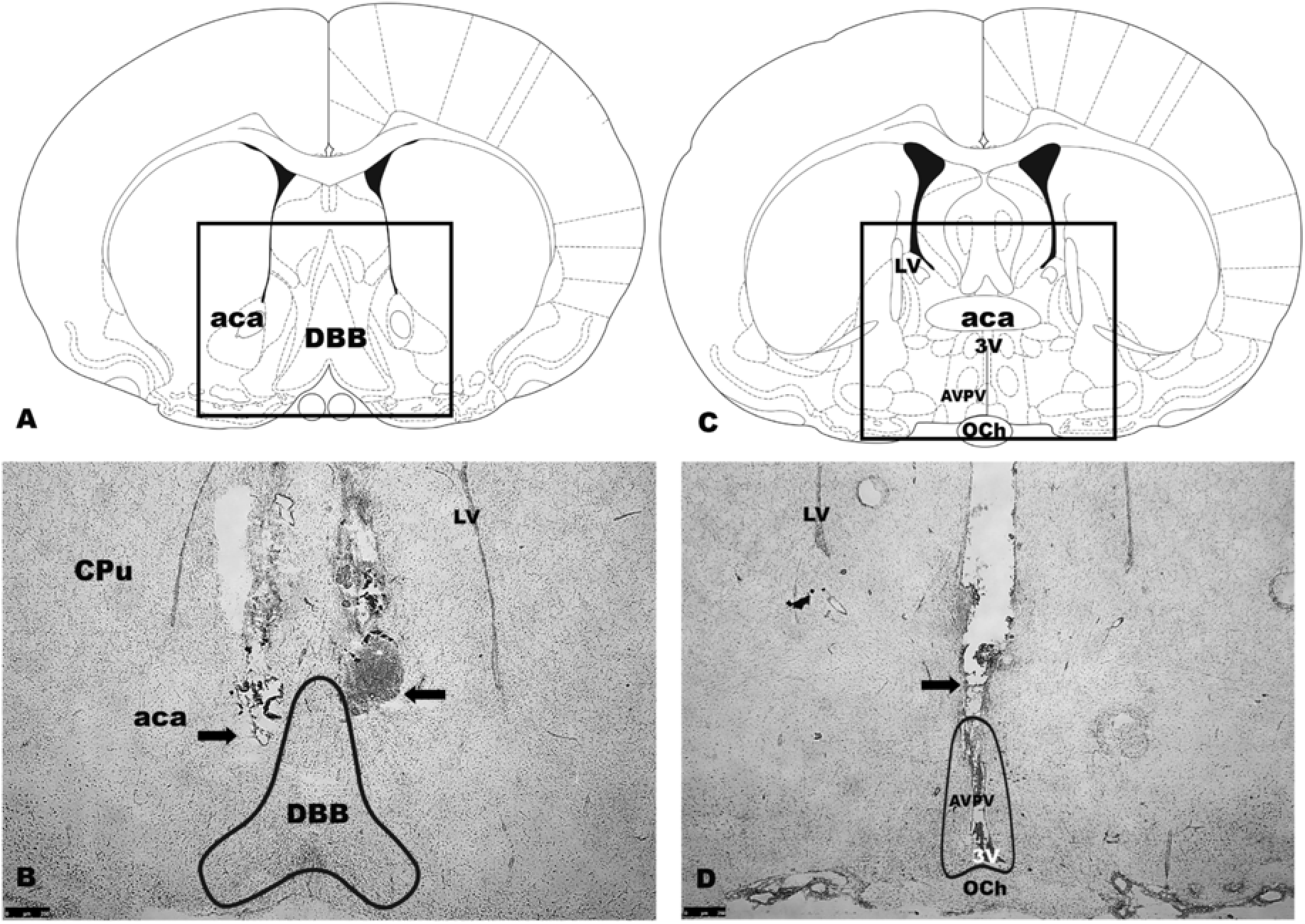
Representative cannula placements for experiments 1 and 2. A) Illustration of sections taken at the level of diagonal band of Broca (DBB; Paxinos & Watson, 2007) and B) bilateral guide cannulae aimed at the DBB in thionin stained coronal section (arrows). C) Illustration of sections taken at the level of anterior ventral periventricular nucleus (AVPV) rostral periventricular region of the third ventricle (RP3V) region (Paxinos & Watson, 2007) and D) unilateral guide cannula aimed at the third ventricle (3V) in thionin stained coronal section (arrow). LV = lateral ventricle; CPu = caudate putamen; aca = anterior commissure, anterior part; OCh = Optic Chiasm. Scale bar = 250 µm.

#### Experiment 2

This experiment tested the subhypothesis that neuroprogesterone-induced kisspeptin release is critical for triggering the LH surge site-specifically in the RP3V. In short, kisspeptin asODNs were infused into the AVPV to reduce EB-induced kisspeptin synthesis and in turn, reduce the releasable pool of kisspeptin, which is expected to block the neuroprogesterone-induced LH surge. On day 13, one week prior to steroid treatment and drug infusion, adult female ovx/adx rats were implanted with a single 3V cannula at the level of the RP3V. On day 20 post ovx/adx surgery, a group of animals (n = 6) were infused with scrambled ODN (scODN) cocktails (10 µg/1 µl) at 1100 hours and were then treated with safflower oil (0.1 ml; s.c.) at 1200 hours. These animals received the same infusion of scODN cocktail at 1100 hours for two subsequent days. Another group of animals (n = 5) received scrambled ODN (scODN) cocktails (10 µg/1 µl) infusion at 1100 hours and were then treated with EB at 1200 hours. These animals received the same infusion of scODN cocktail at 1100 hours on days 21 and 22. A third group of animals (n = 6) were infused on day 20 with antisense ODN (asODN) cocktails (10 µg/1 µl) at 1100 hours and were then treated with oil (0.1 ml; s.c.) at 1200 hours. These animals received the same infusion of asODN cocktail at 1100 hours on days 21 and 22. A fourth group of animals (n=10) were infused on day 20 with asODN cocktails (10 µg/1 µl) at 1100 hours and were then treated with EB at 1200 hours. These animals received the same infusion of asODN cocktail at 1100 hours on days 21 and 22. On Day 22 at 1730 hours, trunk blood and brains were collected approximately 53 hours post EB treatments. Serum LH concentrations were measured by ELISA (Shibaygi via BioVendor; Asheville, NC). Brains were flash-frozen and sectioned at 20 µm and thionin stained for confirmation of cannula placement. Results were analyzed using a one-way ANOVA followed by a post-hoc SNK test.

#### Experiment 3

This experiment was designed to test whether the antagonism of RP3V kisspeptin signaling through its receptor (GPR54) on DBB GnRH neurons blocks the neuroprogesterone-induced LH surge. Initially, we tested whether blocking GPR54 with the kisspeptin receptor antagonist (K-234) also prevents the LH surge. On day 13, one week prior to steroid treatment and drug infusion, adult female ovx/adx rats were implanted with a bilateral cannula aimed at the DBB. At 20 days post ovx/adx surgery, all animals were treated with EB at 1200 hours. On Days 20-22 at 0800 hours, all animals were also given dimethyl sulfoxide (DMSO; 0.1 ml; s.c.). On Day 22 of the experiment, one group of animals (n = 4) underwent two bilateral DBB saline infusions (0.5 µl/per side) at 1500 hours and 1630 hours (Figure 3). A second group of rats (n = 7) received two DBB infusions of K-234 at 1500 hours and 1630 hours on day 22. Approximately 53 hours post EB treatment (1730 hours on Day 22) trunk blood and brains were collected for measurement of serum LH concentration by ELISA and cannula tract placement confirmation, respectively (Figure 3).

The second part of this experiment tested whether K-234 blocked the LH surge in the presence of GPR54 agonist, kisspeptin-10. One week before steroid treatment and drug infusions, adult female ovx/adx Long-Evans rats were implanted with a bilateral cannula aimed at the DBB. At 20 days post ovx/adx surgery, all animals were treated with EB at 1200 hours. On Days 20-22 at 0800 hours, all animals were given AGT (1 mg/kg on Days 20-21 and 5 mg/kg on Day 22; s.c.). On Day 22 of the experiment, one group of animals (n=8) received two DBB infusions of saline (0.5 µl/per side) at 1500 hours and then at1630 hours. A second group of rats (n=6) on day 22 received a DBB infusion of saline at 1500 hours and an infusion of kisspeptin-10 (10 nmol/0.5 µl/per side) at 1630 hours. Importantly, a final group of animals (n=4) received a DBB infusion of K-234 (1 nmol/0.5 µl/per side; Tocris Bioscience; Minneapolis, MN and a gift from Robert P. Millar) at 1500 hours followed by an infusion of kisspeptin-10 (10 nmol/0.5 µl/per side) at 1630 hours. The dose of K-234 was chosen based on a previous study in rats (Grachev et al., 2012).

### Tissue Collection and Cannula Tract Confirmation

For all three experiments, animals were deeply anesthetized with isoflurane and killed by decapitation at 1730 hours on Day 22 - 53 hours post EB treatment. Immediately following decapitation, trunk blood and brains were collected. Blood was allowed to clot for 90 minutes at room temperature and then centrifuged at 2000 x g for 15 minutes at room temperature to separate the serum. Serum was collected and stored at −80° C until analysis for LH by ELISA. Extracted brains were flash frozen on dry ice and stored at −80° C until sectioned for cannula placement or dissected to measure knockdown of kisspeptin mRNA. To visualize cannula placement, brains were mounted onto a chuck using HistoPrep medium, sectioned in the coronal plane at 20 μm through the DBB or 3V/RP3V, and directly mounted onto Superfrost™ Plus slides. Slides were allowed to dry on a slide warmer and stored at −80° C until thionin staining for visualization of cannula tract placement with brightfield microscopy (Figure 2).

### Rat LH Sandwich ELISA

For all experiments, serum LH concentrations were measured by rat LH sandwich ELISA (Shibayagi via BioVendor; Asheville, NC) and performed according to the manufacturer’s protocol. Standards and samples were run in duplicate. Results of the ELISA were measured by a colorimetric microplate reader, VarioSkan 2.2 (ThermoScientific Inc., U.S.A.) at 450 nm. The mean for each sample was calculated and group means were graphed and statistically analyzed. The limits of detection of LH was 0.625 ng/ml. The intra-assay coefficients of variation estimated in two replicated assays of serum samples for experiments 1, 2, and 3 were 7.8%, 2.9%, and 6.8% respectively, and the inter-assay coefficients of variation was 7.8%.

### Western Blot

Brains were harvested on Day 22 of Experiment 2 and kept at −80° C until processing. Blocks containing RP3V and ARH were microdissected from each brain and homogenized in RIPA lysis buffer with protease inhibitors (Santa Cruz Biotechnology; Dallas, TX). Samples were analyzed for protein content using the BCA Spec method (Thermo Scientific Pierce Protein Biology; Waltham, MA). Protein (25 µg) was loaded into each lane of a polyacrylamide gel and electrophoresed. Western blotting was performed using an antibody directed towards rat kisspeptin (1:500; EMD Millipore AB9754; Billerica, MA; Table 2). This antibody has been extensively characterized (Desroziers et al., 2010; Franceschini et al., 2006; Mittelman-Smith et al., 2012). Secondary antibody was applied for one hour (1:10,000 HRP-goat-anti-rabbit; Jackson Immuno; West Grove, PA) and developed using a FluorChemE imager (Protein Simple; San Jose, CA). Blots were stripped and re-probed using an antibody directed towards GAPDH (1:10,000; Millipore MAB374; Billerica, MA) as a loading control. Bands were analyzed by densitometry using AlphaView software (Protein Simple; San Jose, CA). Total signal was achieved by dividing kisspeptin band densitometries by those of GAPDH. The ratio of kisspeptin protein expression to GAPDH was expressed as a percent of the calculated scrambled ODN kisspeptin protein ratio (Figure 5).

**Table 2.** Antibody Table

| **Peptide/**<br>**Protein**<br>**Target** | **Antigen Sequence (If**<br>**Known)** | **Name of Antibody** | **Manufacturer,**<br>**Catalog number,**<br>**and/or Name of**<br>**Individual**<br>**Providing the**<br>**Antibody** | **Species Raised in**<br>**Monoclonal or**<br>**Polyclonal** | **Dilution**<br>**Used** |
| --- | --- | --- | --- | --- | --- |
| Kisspeptin | Mouse kisspeptin-10: YNWNSFGLRY | Anti-Kisspeptin | Millipore,<br>AB 9764 | Rabbit Polyclonal | 1:500 |
| Rabbit IgG |  | HRP-goat-anti-rabbit | Jackson Immuno<br>111-035-047 | Goat | 1:10,000 |
| GAPDH | Rabbit<br>glyceraldehyde-3-phosphate<br>dehydrogenase | 6C5 | Millipore,<br>MAB374 | Mouse<br>Monoclonal | 1:10,000 |
| Mouse IgG |  | HRP-goat-anti-mouse | Jackson immuno<br>111-035-003 | Goat | 1:10,000 |

### Statistical Analysis

Serum LH levels in experiments with three or more groups were analyzed by one-way ANOVA followed by a post-hoc Student-Newman-Keuls (SNK) test (SigmaStat version 3.5). Experiments with only two groups were analyzed by t-test. For western blot experiments, data were analyzed by a one-tailed t-test with unequal variance. For all statistical analyses, values were considered significant at P ≤ 0.05. The ability of kisspeptin K-234 to block an EB-induced LH surge were analyzed using a t-test. Results of the ability of kisspeptin-234 to block the LH surge in the presence of kisspeptin were analyzed using a one-way ANOVA followed by a SNK test.

## RESULTS

### Experiment 1

Inhibiting neuroprogesterone synthesis with AGT blocked EB induction of the LH surge (Figure 4) as indicated by similar LH levels between the Oil + AGT and EB + AGT groups (one-way ANOVA df = 3,32; F = 11.542; P < 0.001; SNK P = 0.384). Subsequent treatment of EB + AGT animals with either systemic progesterone or DBB infusion of kisspeptin-10 stimulated a significant increase in LH compared with either EB + AGT or Oil + AGT saline infused animals (SNK P < 0.05). Both progesterone treatment and DBB kisspeptin-10 infusion produced similar serum LH levels (SNK P = 0.078).

**FIGURE 4.**
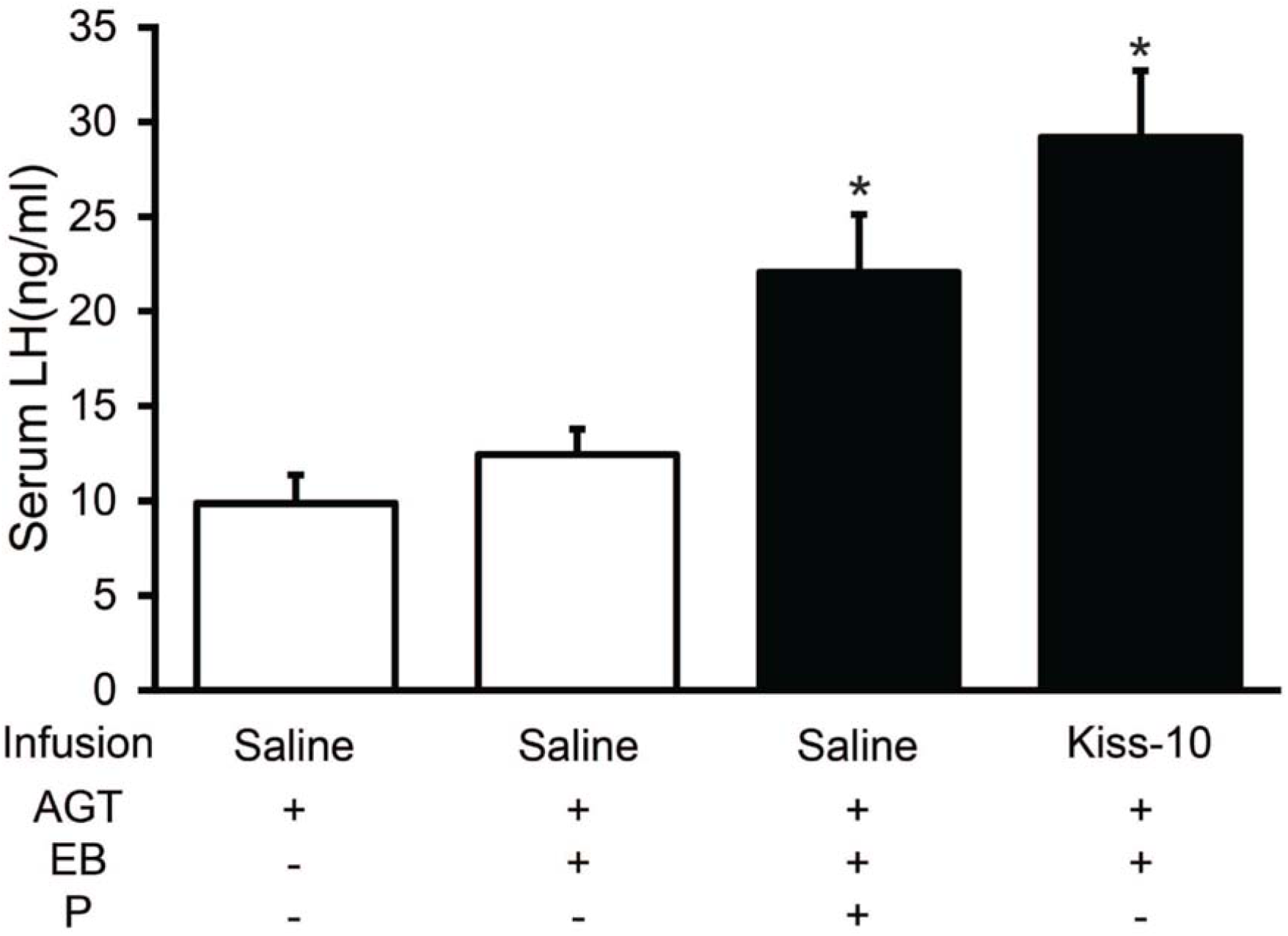
Diagonal band of Broca kisspeptin infusion and subcutaneous progesterone (P) treatment restore the LH surge in ovx/adx rats with inhibited progesterone synthesis. Serum LH measured by ELISA in rats treated with aminoglutethimide (AGT) and either oil or 50 µg EB, and then infused with either saline or kisspeptin-10 (Kiss-10) into the DBB. The group treated with EB + AGT + Kiss-10 had significantly higher levels of LH than both the control groups (Oil + AGT and EB + AGT), but was not significantly different than the positive control group (EB + AGT + P). * indicates significantly greater than Oil + AGT and EB + AGT groups; P < 0.05 (SNK).

### Experiment 2

To test whether kisspeptin synthesized in the RP3V region was necessary for the estradiol-induced LH surge, kisspeptin asODNs infused into the 3V in EB and oil treated animals produced a marked decrease in RP3V kisspeptin protein as assayed by western blot (39.96 ± 18.3% in asODN EB-treated animals versus 99.8 ± 27.4% in scODN EB-treated animals; p = 0.042, t-test, unequal variance). As a verification that ODN did not spread along the 3V, kisspeptin in the ARH was similar in asODN and scODN (controls) treated rats (131.0 ± 73.0% asODN vs. 100.0 ± 15.7% scODN; P = 0.2 in t-test; Figure 5). As expected, LH levels in scODN and kisspeptin asODN rats primed with oil were not significantly different from each other (one-way ANOVA df = 3,26; F = 4.216; P = 0.016; SNK P = 0.852; Figure 6). In contrast, scODN infused animals with EB treatment had significantly increased serum levels of LH compared with oil-primed rats that were infused with kisspeptin asODN or scODN (SNK P = 0.029). Reducing kisspeptin expression with 3V infusion of kisspeptin asODNs significantly reduced LH levels in EB-primed rats compared with scODN treated EB-primed rats (SNK P = 0.011). Importantly, LH levels in kisspeptin asODN + Oil and kisspeptin asODN + EB animals were not significantly different (SNK P = 0.671; Figure 6).

**FIGURE 5.**
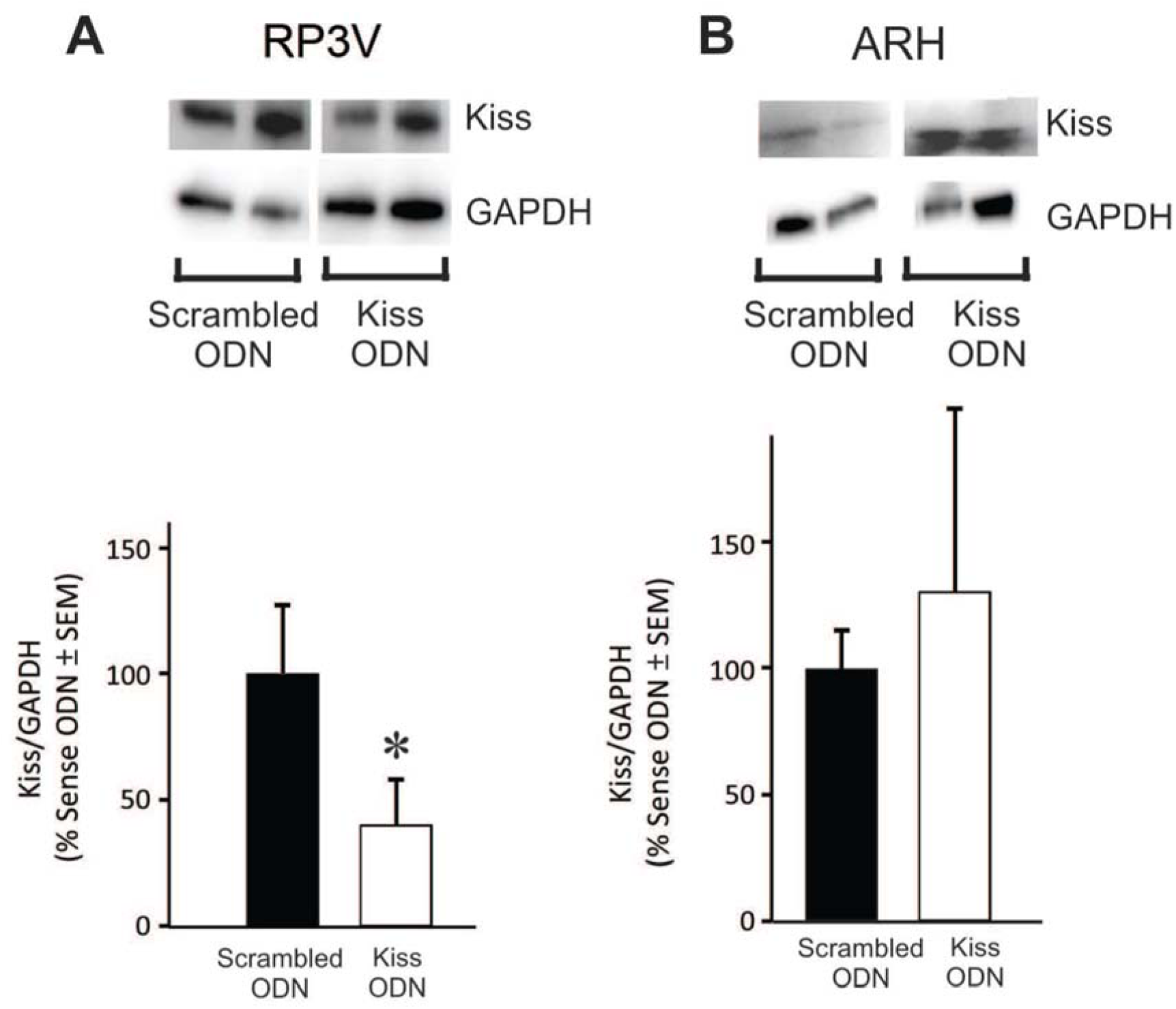
Kisspeptin (Kiss) expression after 3V sense and antisense oligodeoxynucleotide (ODN) infusions in the RP3V (A) and arcuate nucleus (B). A) Representative western blots of Kiss protein expression and GAPDH in the RP3V region of EB primed rats infused with either scrambled ODNs or kisspeptin ODNs (Kiss ODN). Histogram illustrates the ratio of Kiss protein expression to GAPDH expression as a percent of scrambled ODN. B) Representative western blots of Kiss protein expression and GAPDH in the arcuate nucleus of the hypothalamus of EB primed rats infused with either scrambled ODNs or kisspeptin ODNs (Kiss ODN). Histogram illustrates the ratio of Kiss protein expression to GAPDH expression as a percent of scrambled ODN. * indicates less than scrambled ODN; P < 0.05.

**FIGURE 6.**
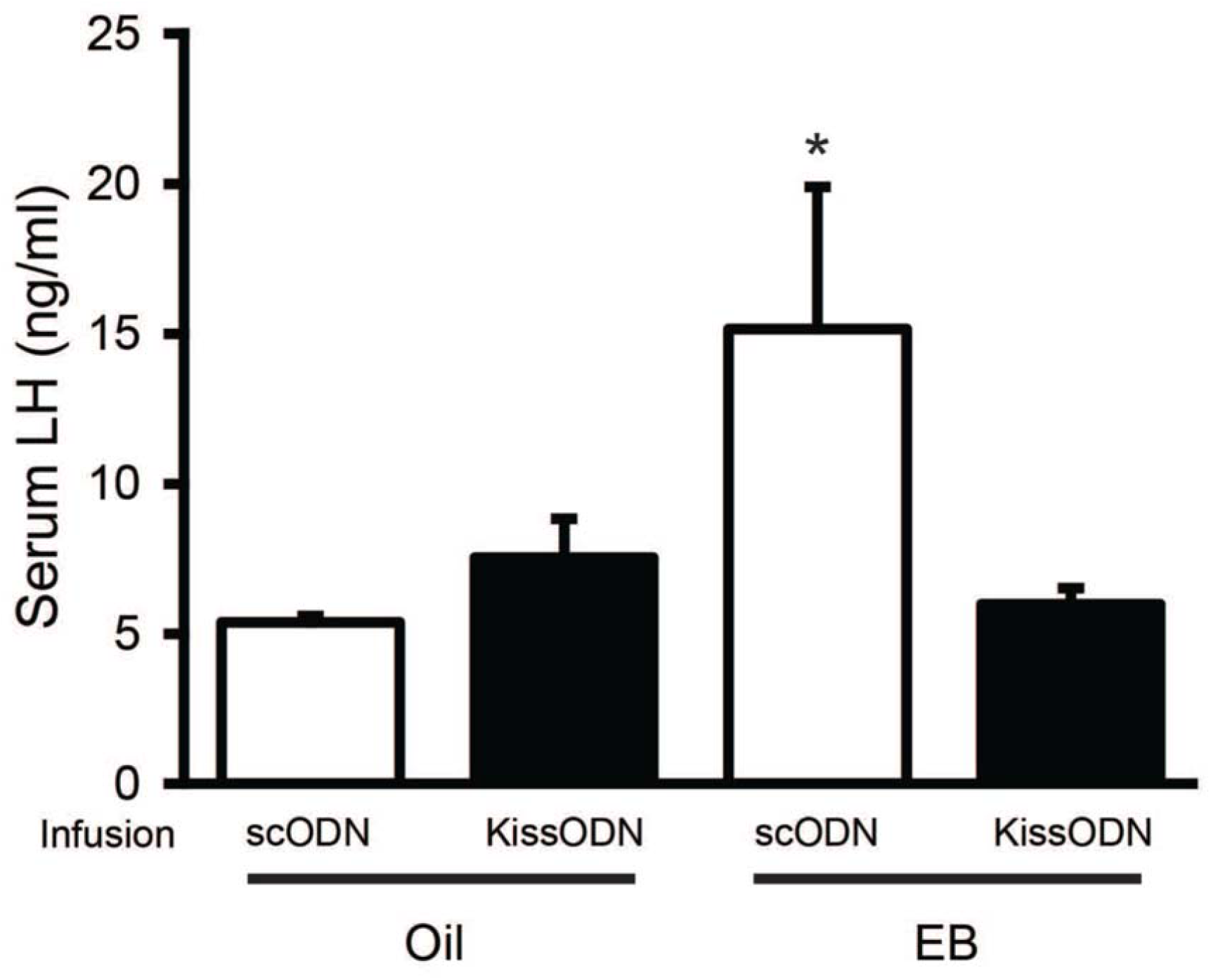
RP3V kisspeptin antisense ODN (KissODN) cocktail infusion blocked EB induction of the LH surge in ovx/adx rats. Serum LH levels were measured by ELISA in rats treated with either oil or 50 µg EB and infused with either KissODN or scrambled (scODN) into the 3V at the level of the RP3V. scODN + Oil and KissODN + Oil controls both displayed significantly lower serum LH levels than the positive control group, scODN + EB. Importantly, KissODN + EB-treated animals also had significantly lower mean serum LH concentration than scODN + EB-treated animals, and KissODN + Oil- and KissODN + EB-treated animals did not have significantly different levels of serum LH. * indicates significantly greater than other groups P < 0.05 (SNK).

### Experiment 3

To determine whether neuroprogesterone induced kisspeptin release activated the kisspeptin receptor, GPR54, to trigger the LH surge, GPR54 was antagonized by infusion of K234 into the DBB. In EB-primed rats, infusion of K-234 significantly reduced serum LH levels compared with saline infused animals (t-test df = 9; t = 2.439; P = 0.037; Figure 7). AGT administration that blocked the increase in LH release in EB-primed rats was reversed by kisspeptin infusion into the DBB, as evidenced by significantly increased LH levels (one-way ANOVA df = 2,17; F = 6.575; P = 0.009; SNK P = 0.008). Infusion of kisspeptin-234 into the DBB also blocked the kisspeptin induced increase in LH compared with saline infused animals (SNK P = 0.026; Figure 8). Importantly, no significant difference in serum LH levels was observed between EB + AGT + Saline treated animals and EB + AGT + kisspeptin-234 + kisspeptin-10 treated animals (SNK, P = 0.634).

**FIGURE 7.**
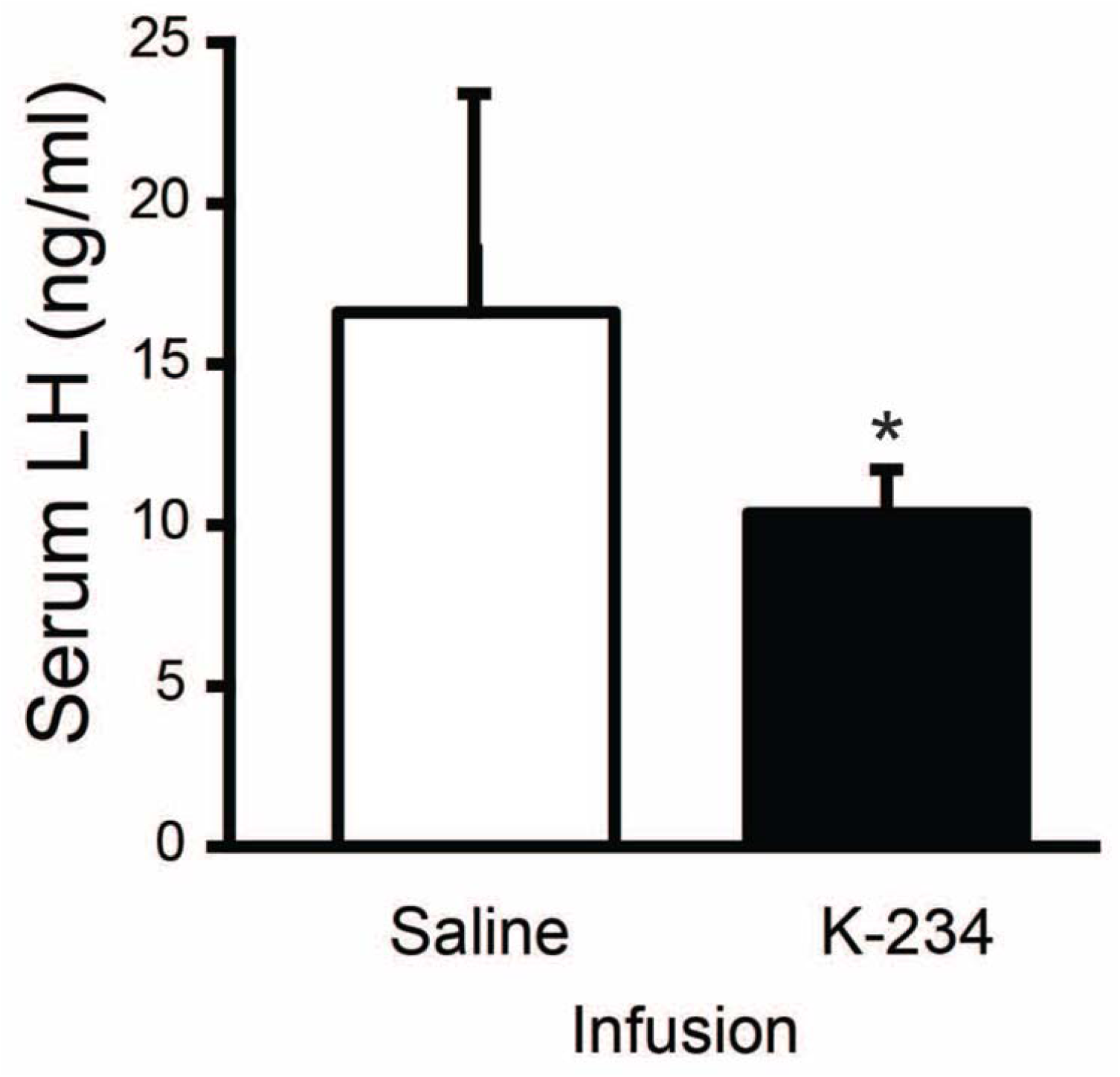
Diagonal band of Broca infusion of the GPR54 receptor antagonist, kisspeptin-234 (K-234), blocks EB induction of the LH surge in ovx/adx rats. Serum LH concentration was measured by ELISA in rats treated with 50 µg EB and DMSO and infused with either saline or K-234. K-234-infused animals had significantly reduced mean serum LH levels compared to saline-infused control animals. * indicates significantly less than other group; P < 0.05.

**FIGURE 8.**
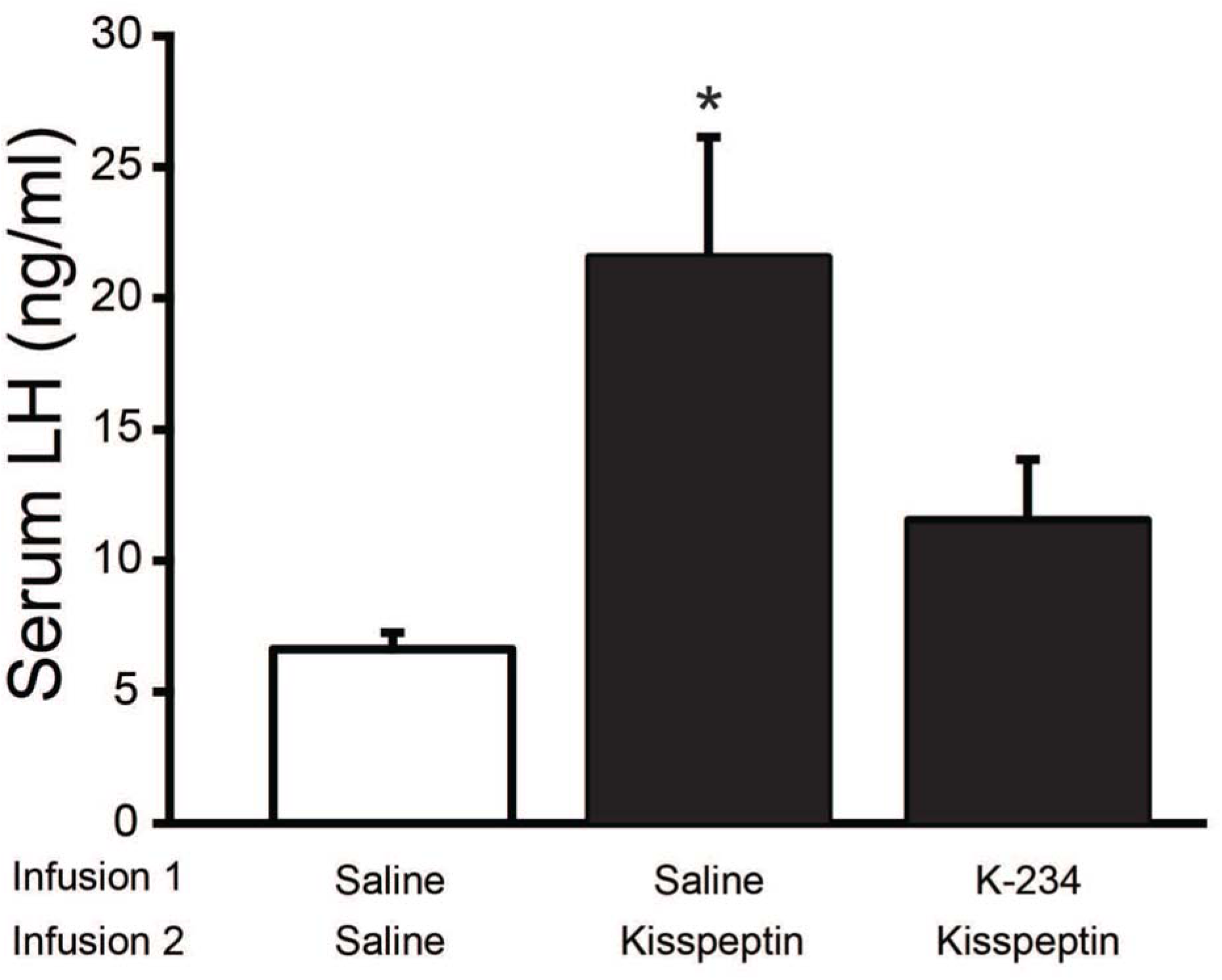
Diagonal band of Broca infusion of the GPR54 receptor antagonist, kisspeptin-234 (K-234), blocked the EB- and kisspeptin-induced LH surge in EB + AGT treated ovx/adx rats. Serum LH levels were measured by ELISA in animals treated with 50 µg EB, the progesterone synthesis inhibitor, AGT (1 and 5 mg/kg) and then given a DBB infusion of either saline or K-234 (2 nmol/ 1 µl), followed by a second infusion of the GPR54 agonist, Kiss-10 (20 nmol/1µl), 15 minutes later. EB + AGT + K-234 + Kiss-10 treated animals had significantly reduced serum LH levels when compared to EB + AGT + Saline + Kiss-10 treated animals, and was not significantly different from EB + AGT + Saline treated control animals. * indicates significantly greater than other groups; P < 0.05 (SNK).

## DISCUSSION

The major finding of these experiments is that estradiol-induced neuroprogesterone triggers the LH surge via activation of RP3V kisspeptin neurons in the DBB - a result congruent with the idea that locally synthesized neuroprogesterone activated kisspeptin neurons to stimulate GnRH neurons inducing the LH surge. In estradiol-primed, ovx/adx rats with AGT-blocked neuroprogesterone synthesis (P. Micevych et al., 2003), treatment with progesterone or site-specific infusions of kisspeptin into the DBB produced a LH surge. Inhibiting kisspeptin synthesis in the RP3V with asODN blocked the estradiol/neuroprogesterone induction of the LH surge. Further, infusion of the KissR1R (GPR54) antagonist, K-234, blocked estradiol- or kisspeptin-induced LH surges confirming that the release of kisspeptin is downstream of estradiol induced neuroprogesterone synthesis for the LH surge.

Based on in vitro studies, we had hypothesized that kisspeptin neurons serve as the site of integration of temporal steroid hormone actions that results in kisspeptin release that activates DBB GnRH neurons to trigger the LH surge (Figure 1; (P. Micevych & Sinchak, 2008b; Mittelman-Smith et al., 2015; Mittelman-Smith et al., 2018). It is likely that neuroprogesterone acts on estradiol-induced PGR in RP3V kisspeptin neurons to induce kisspeptin release. Both RP3V kisspeptin neurons *in vivo* (Clarkson et al., 2008; Smith et al., 2005; Smith, Popa, Clifton, Hoffman, & Steiner, 2006) and mHypoA51 immortalized kisspeptin neurons (Mittelman-Smith et al., 2015) express ERα and PGR (Clarkson et al., 2008; Mittelman-Smith et al., 2015; Mittelman-Smith et al., 2018; Smith et al., 2006). *In vitro*, the mHypoA51 neurons express ERα intracellularly and on the membrane. Activation of nuclear ER is needed for induction of PR. However, a portion of ERα which is trafficked to the membrane and when activated increases the free cytoplasmic calcium. *In vitro* studies using mHypoA51 cells support these findings (Mittelman-Smith, Scott, Wong, & Micevych, 2016; Mittelman-Smith et al., 2018). When co-cultured with post-pubertal female hypothalamic astrocytes, estradiol treatment results in a significant increase in kisspeptin protein expression (Mittelman-Smith et al., 2018). Subsequently, *in vivo*, this kisspeptin is released via induction of astrocytic neuroprogesterone synthesis, as evidenced by the progesterone induction of kisspeptin release (Mittelman-Smith et al., 2018). Estradiol induces kisspeptin release in co-cultures with primary astrocytes (the source of estradiol-induced neuroprogesterone), suggesting that progesterone-induced kisspeptin release is occurring directly through kisspeptin neurons themselves (Mittelman-Smith et al., 2018). While we cannot yet definitively exclude the possibility that one or more other interposed neurons are involved in this circuit, the present data strongly indicate that neuroprogesterone directly activates RP3V kisspeptin neurons to trigger the LH surge, and is supported by the *in vitro* results with mHypoA51 cells (Mittelman-Smith et al., 2018).

*In vivo*, we have previously demonstrated that inhibiting progesterone synthesis with AGT blocks the EB-induced LH surge in ovx/adx rats (P. Micevych et al., 2003), and AGT specifically in the hypothalamus arrested intact cycling rats in a proestrous state prior to the LH surge (P. Micevych & Sinchak, 2008a). Here, we rescued the LH surge in AGT + EB-treated rats by kisspeptin infusion into the DBB (Figure 4). Conversely, the EB-induced LH surge was blocked by DBB infusion of K-234 (Figure 7), which demonstrates that the actions of kisspeptin are downstream of estradiol and neuroprogesterone. These findings support the idea that neuroprogesterone induces the release of kisspeptin to trigger the LH surge.

The present study also demonstrates that EB-treatment induces kisspeptin synthesis important for induction of the LH surge (Figure 5 & 6). To date, experiments have not addressed whether kisspeptin is induced only by estradiol *in vivo* because estradiol also stimulates neuroprogesterone synthesis, which can also augment kisspeptin expression. *In vitro*, estradiol induces kisspeptin mRNA and protein (Mittelman-Smith et al., 2015) but neuroprogesterone augments kisspeptin expression and mobilizes calcium to enhances kisspeptin release needed for activating GnRH neurons and the LH surge (Mittelman-Smith et al., 2015; Mittelman-Smith et al., 2018). The present results show that kisspeptin asODNs blunted RP3V kisspeptin protein expression and successfully blocked the EB-induced LH surge, which are congruent this idea (Figure 5 & 6). Further, the current results support the idea that kisspeptin produced specifically in the region of the RP3V (as opposed to kisspeptin in the ARH) is required for GnRH neuron activation for subsequent induction of the LH surge since the infusions of kisspeptin asODNs did not have an effect on kisspeptin expression in the ARH (Figure 5). Kisspeptin neurons originating in the RP3V project to GnRH neurons (Wintermantel et al., 2006) and are responsible for the positive feedback of LH, which results in the LH surge. On the other hand, the population of kisspeptin neurons in the ARH regulates the negative feedback of LH, generating LH pulsatility (Adachi et al., 2007; Kelly, Zhang, Qiu, & Ronnekleiv, 2013; Mittelman-Smith et al., 2012; Smith et al., 2006). Kisspeptin antisense infused directly into the ARH or RP3V regions support our findings. ARH infusions reduced pulsatility and prolonged the estrous cycle, and attenuated but did not blocked spontaneous LH surges. In contrast, RP3V infusions had no effect on pulsatility, but blocked LH surges and prolonged estrous cycles (Beale et al., 2014; Hu et al., 2015). The present studies are consistent with estradiol positive feedback on GnRH neurons via RP3V kisspeptin neurons. Moreover, estradiol regulates the ARH kisspeptin neuron population in a different direction. While estradiol increases kisspeptin expression in the RP3V, it suppresses expression in the ARH (Smith et al., 2005; Smith et al., 2006). Thus, the kisspeptin expression is low in the EB-treated ARH and further knockdown with asODN would be expected to have a measurable physiological impact. This was supported by the observation that local antagonism of the KISS1R in the DBB blocks the LH surge, indicating that RP3V kisspeptin neurons are the primary population responsible for mediating the LH surge.

We do not have a timeline of the estradiol-induced neuroprogesterone/kisspeptin activation of the LH surge in vivo. In vitro, using mHypoA51, immortalized kisspeptin neurons, estradiol induced kisspeptin mRNA within 24 hrs and the protein by 48 hrs (Mittelman-Smith et al., 2015). This matches results from whole animal studies where during the beginning of the cycle, while estradiol negative feedback predominates, estradiol appears to be simultaneously priming the positive feedback circuits by upregulating PGR and kisspeptin expression in RP3V neurons. As the estrous cycle progresses and estradiol positive feedback predominates, the number of RP3V neurons that express kisspeptin mRNA increases (Smith et al., 2006). This produces a releasable pool of kisspeptin peptide whose release is stimulated by neuroprogesterone. Indeed, co-culturing astrocytes with mHypoA51 neurons, we observed that estradiol-induced neuroprogesterone augments kisspeptin release (Mittelman-Smith et al., 2018). In the present experiment, we infused kisspeptin asODN over three days overlapping with EB treatment, which blocked kisspeptin expression and prevented the LH surge.

Neuroprogesterone synthesis in astrocytes is stimulated through membrane-initiated estradiol signaling (Kuo, Hamid, Bondar, Prossnitz, & Micevych, 2010; Kuo, Hariri, Bondar, Ogi, & Micevych, 2009)., releasing internal calcium stores, activating PKA which in turn activates cholesterol carrier proteins stimulating the transport of cholesterol into the mitochondrion where it is converted into pregnenolone (Figure 1; Chen et al., 2014; Kuo, Hamid, Bondar, Dewing, et al., 2010; Kuo, Hamid, Bondar, Prossnitz, et al., 2010; P.E. Micevych et al., 2007; Sinchak et al., 2003). Similarly, in mHypoA51 neurons, estradiol induces kisspeptin mRNA and protein through a membrane initiated action, which was demonstrated by the efficacy of membrane-impermeable 17β-estradiol conjugated to BSA (E-BSA) to induce kisspeptin expression (Mittelman-Smith et al., 2015). In the mHypoA51 cells, E-BSA did not induce PGR expression, which was readily induced by free estradiol, suggesting a direct nuclear action in line with previous results (Mittelman-Smith et al., 2015).

The findings reported in the present study support the neuroprogesterone-LH surge model, and reinforce the concept of estradiol initiation and regulation of the synchronization of the events occurring in the brain with ovulation. In gonadally intact rats, positive feedback of estradiol is essential for triggering the LH surge, while in ovx/adx rats, exogenous EB is sufficient to trigger a physiologically comparable LH surge (P. Micevych et al., 2003). In the beginning of the estrous cycle (diestrus I and II), the LH surge circuit is under estradiol negative feedback, and female rats are not sexually receptive (reviewed in Blaustein & Mani, 2007). Simultaneously, estradiol acts on kisspeptin-producing neurons in the RP3V, resulting in increased synthesis and expression of PGR and kisspeptin (Kinoshita et al., 2005; Smith et al., 2006; Wintermantel et al., 2006). As estradiol peaks, it stimulates hypothalamic astrocytes to upregulate the expression and activity of the progesterone synthesis enzyme, 3β-hydroxysteroid dehydrogenase, resulting in the synthesis and release of neuroprogesterone (Kuo & Micevych, 2012; Soma, Sinchak, Lakhter, Schlinger, & Micevych, 2005) prior to the LH surge. This neuroprogesterone then acts on PGR-expressing kisspeptin neurons in the RP3V to release kisspeptin, stimulating GPR54 on GnRH neurons. Activation of GPR54 induces the secretion of GnRH into the hypothalamo-hypophyseal portal system causing the surge release of LH from the anterior pituitary. Therefore, the switch from estrogen negative to positive feedback appears to be two-fold, involving: i) estradiol-induced upregulation of kisspeptin and PGR in kisspeptin neurons, and ii) estradiol-induced synthesis of astrocyte-derived neuroprogesterone.

In summary, these data are consistent with the hypothesis that ovarian estradiol appears to synchronize the synthesis of PGR, kisspeptin and neuroprogesterone, which activates RP3V neurons to release kisspeptin activating GnRH neurons inducing the LH surge. Together, these findings support the idea that an important step in regulating ovulation and luteinization is the interaction of ovarian estradiol and hypothalamic astrocytes derived neuroprogesterone to induce and release kisspeptin.

## Acknowledgements

Authors thank Dream Le, Lam Nguyen, Karla Romero, Rachael Bonifacio, Amelia Welborn, Jessica Phan, Chhorvann Serey, Maribel Maciel, Daniel Tran and Lauren Jewel for their technical assistance and Emo Gonzales for his assistance on illustrations. Research was
supported by NIH grant HD042635 (PM) HD058638 (KS) 5R25GM071638 (NIH RISE–CSULB), 5T32HD007228 (MMS).

